# Endothelial Cells are Heterogeneous in Different Brain Regions and are Dramatically Altered in Alzheimer’s Disease

**DOI:** 10.1101/2023.02.16.528825

**Authors:** Annie Bryant, Zhaozhi Li, Rojashree Jayakumar, Alberto Serrano-Pozo, Benjamin Woost, Miwei Hu, Maya E. Woodbury, Astrid Wachter, Gen Lin, Taekyung Kwon, Robert V. Talanian, Knut Biber, Eric H. Karran, Bradley T. Hyman, Sudeshna Das, Rachel Bennett

## Abstract

Vascular endothelial cells play an important role in maintaining brain health, but their contribution to Alzheimer’s disease (AD) is obscured by limited understanding of the cellular heterogeneity in normal aged brain and in disease. To address this, we performed single nucleus RNAseq on tissue from 32 AD and non-AD donors each with five cortical regions: entorhinal cortex, inferior temporal gyrus, prefrontal cortex, visual association cortex and primary visual cortex. Analysis of 51,586 endothelial cells revealed unique gene expression patterns across the five regions in non-AD donors. Alzheimer’s brain endothelial cells were characterized by upregulated protein folding genes and distinct transcriptomic differences in response to amyloid beta plaques and cerebral amyloid angiopathy (CAA). This dataset demonstrates previously unrecognized regional heterogeneity in the endothelial cell transcriptome in both aged non-AD and AD brain.

**Significance Statement:** In this work, we show that vascular endothelial cells collected from five different brain regions display surprising variability in gene expression. In the presence of Alzheimer’s disease pathology, endothelial cell gene expression is dramatically altered with clear differences in regional and temporal changes. These findings help explain why certain brain regions appear to differ in susceptibility to disease-related vascular remodeling events that may impact blood flow.

## Introduction

Cerebral vasculature plays a central role in the maintenance of brain health, and in Alzheimer’s disease (AD), changes that impact blood flow and blood-brain barrier function may exacerbate neurodegeneration and cognitive decline (Iturria-Medina et al., 2016; Montagne et al., 2016; Benedictus et al., 2017; Leijenaar et al., 2017; Nation et al., 2019). A growing body of literature suggests that diverse blood vessel changes including aberrant angiogenesis, vascular pruning, inflammation, senescence, and other remodeling events occur alongside AD-related pathological aggregation of amyloid beta and tau (Brown and Thore, 2011; Bennett et al., 2018; Bryant et al., 2020; Lau et al., 2020). However, our current understanding is limited by the lack of characterization of the regional heterogeneity observed in the human brain, particularly in non-neuronal cell-types such as endothelial cells. Indeed, few human studies have deeply characterized vasculature from multiple brain regions and comparisons with mouse models may be inadequate (Hodge et al., 2019). Thus we have, at best, an incomplete picture of how these vascular mechanisms underlie functional alterations that affect disease progression.

Previous work has highlighted the heterogeneity of the endothelial transcriptome and translatome between the brain and other organs such as lung, heart, liver, and kidney (Nolan et al., 2013; Jambusaria et al., 2020; Kalucka et al., 2020; Paik et al., 2020). Moreover, advances in single-cell sequencing technologies have enabled the resolution of brain endothelial cell subtypes (Vanlandewijck et al., 2018; Kalucka et al., 2020; Yang et al., 2022). However, such studies have only examined either a singular brain region or a pooled cortical homogenate, which may obscure cerebrovascular heterogeneity at the region-specific level (Bernier et al., 2021). Other groups have recently demonstrated region-specific cellular subtypes in murine astrocytes (Batiuk et al., 2020; Hasel et al., 2021) and microglia (Tan et al., 2019). We reasoned that the endothelial cell transcriptome may also vary by location in the brain, given both developmental and morphological differences in the microvasculature in different brain regions.

Understanding regional heterogeneity in the human brain is important in AD because amyloid beta, neurofibrillary tangles and neurodegeneration do not affect all brain regions to the same extent. Numerous neuropathological studies have outlined the spatial distributions of these changes. Amyloid deposits in region-specific patterns are distinct for plaques and for cerebral amyloid angiography (CAA). The temporal and frontal association cortices accumulate more plaques than primary sensory areas such as visual cortex (Haroutunian et al., 1998; Serrano-Pozo et al., 2011) whereas CAA has much higher prevalence in parietal and occipital cortices(Vinters and Gilbert, 1983). Neurofibrillary tangles show yet another distinct hierarchical regional pattern. Braak staging describes the progressive severity of tau neurofibrillary tangle pathology, with early stages affecting primarily the entorhinal cortex (I-II), then limbic structures including the hippocampus (III-IV), followed by spread to isocortical areas including the frontal cortex and association visual cortex regions (V2) before appearing in primary sensory areas (VI) including visual cortex (Braak and Braak, 1991a; Braak et al., 2006a). Notably, neuron loss correlates better with neurofibrillary tangle burden than with amyloid beta plaque burden (Arriagada et al., 1992; Gómez-Isla et al., 1997). For this reason, we focused these studies on cortical regions included in the Braak staging scheme that represent the hierarchical accumulation of tau pathology in AD, in order to examine the effect of disease progression on the endothelial cell population.

Considering that cortical brain areas are differently affected by amyloid beta and tau pathologies, and that vascular changes are intimately related to AD, the goals of this work were two-fold: 1) to characterize baseline heterogeneity in the endothelial cell transcriptome in the normal aged brain; and 2) to elucidate how regional differences in disease progression impact endothelial cell gene expression in AD.

## Methods

### Donor Tissue Collection

Thirty-two donors were selected from the Neuropathology Core of the Massachusetts Alzheimer’s Disease Research Center, which obtained written informed consent from next-of-kin for brain donation. Tissues were assessed by a neuropathologist for AD-related neuropathology burden according to NIA-AA guidelines (Hyman et al., 2012) (Supplementary Table 1; Figure S1) and included high pathology Braak V/VI donors (n=16; B3), intermediate pathology Braak III/IV donors (n=5; B2) and low pathology Braak 0-II donors (n=11; B0/1). For each donor, frozen postmortem cortical tissue was obtained from the following five brain regions where available: entorhinal cortex (EC), inferior temporal gyrus (ITG), prefrontal cortex (PFC), secondary visual cortex (V2), and primary visual cortex (V1). Formalin-fixed, paraffin embedded tissue blocks from the opposite hemisphere were also obtained for histological studies.

### Histology

For validation of targets identified by transcriptomic analysis, paraffin-embedded EC and V1/V2 were sliced on a microtome at 8-microns, and sections were mounted on Superfrost slides. Two sets of sections were prepared, one containing hippocampus, entorhinal cortex, and inferior temporal gyrus at the level of the lateral geniculate nucleus, and the other containing occipital cortex including primary and association visual areas. Sections were rehydrated in xylene and a descending concentration of ethanol. Epitope retrieval was carried out in citrate buffer (pH 6.0, containing 0.05% Tween-20) with microwave heating at 95 °C for 20 min. Sections were then blocked with 5% bovine serum albumin in Tris buffered saline containing 0.25% Triton-X (TBS-X) for one h at room temperature and then incubated with primary antibodies overnight at 4 °C. Primary antibodies used in this study targeted CREM (Invitrogen cat no. PIPA581971), HSPH1 (Proteintech, cat no. 13383-1-AP), HSP90AA1 (Invitrogen cat no. PA3-013), IGFBP3 (Invitrogen cat no. PA5-27190), MT2A (Invitrogen cat no. PA5-102549) and GLUT1 (EMD Millipore cat no. 07-1401). The following day, sections were rinsed three times with TBS and then incubated in Alexa dye conjugated secondary antibodies in TBS-X for 1 h at room temperature. Afterwards, sections were rinsed three times with TBS and Trueblack (Biotum) was applied for 30 s to quench autofluorescence. Sections were rinsed with H2O and then coverslipped with Fluoromount G with DAPI. An Olympus VS120 slide scanner was used to capture digital images using a 20x lens.

For measurement of plaque burden in tissues, cryosections (10 µm thickness) adjacent to those used for single-nucleus (sn)RNAseq were subjected to immunohistochemistry with mouse monoclonal anti N-terminal abeta antibody clone 3D6. The staining was performed on a Leica BOND Rx automated stainer using DAB-based detection (Leica). Sections were scanned in a slide scanner (3DHistech, Pannoramic 250) and area fraction (i.e., % area of tissue section occupied by 3D6 -immunoreactive plaques) was measured using HALO software (Indica Labs, Albuquerque, NM, USA).

For assessing CAA, tissue sections were processed for amyloid beta (clone 6F/3D, Agilent cat no. M0872) DAB histology by the Massachusetts ADRC. Four tissue blocks were assessed which encompassed all five brain regions; V1 and V2 were located in the same block and received the same score. Slides were digitized and evaluated by a blind researcher (B.W.) using a 0-4 CAA grading scheme (Olichney et al., 1995). An average score for each donor was used for analysis.

### ELISA

For quantification of phospho(T231)tau and total tau, 25 mg of tissue was dissected prior to nuclei isolation. Tissue was dounce homogenized with 5 v/w of PBS and then centrifuged to pellet insoluble material at 3,000 x *g* for 5 minutes at 4 °C. The resulting supernatant was reserved for analysis using MesoScale Discovery Kit #K15121D according to the manufacturer’s instructions.

### Isolation of nuclei from postmortem frozen brain tissue

Nuclei isolation and sorting was performed as previously described with minor modifications(Gerrits et al., 2021). Briefly, fresh frozen tissue was cryosectioned (approximately 40 sections of 40 µm thickness each) and lysed in sucrose lysis buffer (10 mM Tris–HCl (pH 8.0); 320 mM sucrose; 5 mM CaCl2; 3 μM Mg(Ac)2; 0.1 mM EDTA; 1 mM dithiothreitol (DTT) and 0.1% Triton X-100). Lysates were filtered through a 70 µm cell strainer. Nuclei were purified by ultracentrifugation (107,000 x *g* for 1.5 h at 4 °C) through a sucrose cushion (10 mM Tris–HCl (pH 8.0); 1.8 M sucrose; 3 μM Mg(Ac)2; 0.1 mM EDTA and 1 mM DTT). Supernatants were removed and pellets were re-suspended in 2% BSA/PBS containing RNase inhibitor (0.2 U/μL) (Roche). Nuclei were incubated with fluorescently conjugated antibodies against the neuronal marker NEUN (RBFOX3/NEUN (1B7) AF647 mouse mAB, Novus Biologicals, NBP1-92693AF647) and the oligodendrocyte transcription factor OLIG2 (Anti-OLIG2 clone 211F1.1 mouse mAb, Merck Millipore, MABN50A4). Samples were kept on ice throughout the isolation and staining procedure. Nuclei were stained with Sytox blue (Thermo Fisher) and sorted on a BD FACSAria Fusion. For each sample, we collected Sytox^pos^ NeuN^pos^Olig2^neg^ and Sytox^pos^NeuN^neg^Olig2^neg^ nuclei. This second group, depleted for neurons and oligodendrocytes, was examined in this study.

### snRNAseq library preparation

Single-nucleus cDNA libraries were constructed using the Chromium Single Cell 3’ Reagents Kit V3. Samples were pooled and sequenced targeting at least 30k reads per cell on an Illumina NovaSeq.

### snRNAseq data preprocessing and quality control

Sytox^pos^NeuN^neg^Olig2^neg^ nuclei raw data was processed with 10X Genomics cellranger software v4, based on GRCh38 pre-mRNA reference and QC’ ed to identify potential sample outliers. Subsequently, raw counts were filtered per region to include only cells with > 100 exonic counts, and > 800 features and counts (both intronic and exonic) as well as a mitochondrial gene percentage < 15%. A sample specific subsequent filter removed additional potential low-quality cells, only including cells higher/lower than the median +/-3*MAD (median absolute deviation) of gene and UMI count numbers per sample. This yielded final matrices of 231085 (ITG) + 202514 (EC) + 280665 (PFC) + 253403 (V1) + 234567(V2) nuclei and 35423 features.

UMI counts were normalized and integrated across donors for each brain region separately, following the recommended Seurat integrative workflow (Stuart et al., 2019). For each brain region, the filtered count matrices were log-normalized and the top 1,000 highly variable genes (HVGs) were identified for each donor using FindVariableFeatures() with selection.method = “vst” Anchoring features for cross-sample integration were identified using FindIntegrationAnchors() with reduction= “rpca” for reciprocal principal component analysis (rPCA), as is recommended for larger datasets (see https://satijalab.org/seurat/articles/integration_rpca.html). The total number of reads (nCount_RNA) and percent mitochondrial reads (percent.mito) was regressed out. Principal component analysis (PCA) was applied for dimensionality reduction using RunPCA() with npcs=50. The PCA-reduced dataset was further passed to uniform manifold approximation and projection (UMAP) for nonlinear dimensionality reduction using the RunUMAP() function, with dims=1:30. Graph-based shared nearest neighbor (SNN) clustering was applied using FindClusters() with resolutions 0.1 through 1. Subcluster resolution 0.1 was selected to identify clusters of the following cell types: astrocyte, microglia, neuron, endothelial cell, oligodendrocyte, or other. Cell types were manually identified using DimPlot and heatmap visualization of canonical cell type markers (see Supplementary Table 2 for markers used).

The Seurat objects for each brain region were subsetted to only the clusters identified as endothelial cells (Figure S2B), and the endothelial subclusters were re-integrated using canonical correlation analysis (CCA) within each brain region. Specifically, anchor features were identified based on the top 2,000 HVGs and were passed to FindIntegrationAnchors() with k.filter=20. The data were CCA-integrated using IntegrateData() with k.weight=20 and scaled using ScaleData(), with the total number of reads (nCount_RNA) and percent mitochondrial reads (percent.mito) regressed out. PCA was applied for dimensionality reduction using RunPCA() with npcs=50. The resulting scree plot was visualized to select the number of components to retain for graph-based clustering for each brain region (Figure S3A). We chose the following numbers of components: EC, 8; ITG, 9; PFC, 7; V2, 8; V1, 9. UMAP dimensionality reduction and SNN clustering were performed as described above. The final region-wise endothelial cell matrices for each brain region had the following cell counts: EC, 4,271; ITG, 10,850; PFC, 12,638; V2, 15,553; V1, 18,235.

These endothelial matrices were then rPCA-integrated across brain regions using FindIntegrationAnchors() with the top 2,000 HVGs and IntegrateData() with k.weight=20. The inter-region integrated dataset was scaled using ScaleData() with nCount_RNA and percent.mito as variables to regress out. Dimensionality reduction was performed using RunPCA() with npcs=50 and and RunUMAP() with dims=1:30. Graph-based shared nearest neighbor (SNN) clustering was applied using FindClusters() with resolutions 0.1 through 1. Residual contamination from non-endothelial cell types was evaluated using the DoHeatmap() function with known cell type markers (Supplementary Table 2) using subcluster resolution 0.2 (Figure S3B). Subclusters 4, 5, 7, and 10 were removed based on upregulation of genes related to one or more non-endothelial cell types and/or general downregulation of endothelial marker genes.

### Vessel segment identification

In order to examine AD-related changes along the cerebral arteriovenous axis (Vanlandewijck et al., 2018; Chen et al., 2020; Zhao et al., 2020), we used endothelial zonation-specific gene markers from a recent large-scale brain endothelial sequencing study (Yang et al., 2022) (Supplementary Table 3). This set included 62 arterial markers, 129 capillary markers, and 87 venous markers. For each donor and brain region, we tabulated the geometric mean of the log-normalized expression across the arterial, capillary, and venous gene markers, respectively. Each endothelial nucleus was classified according to the segment with the largest geometric mean expression of the three types. Sequencing characteristics for each vessel segment are depicted in Figure S4A-B.

### Differential expression analysis

All differential expression (DE) analysis was performed using either the FindMarkers() or the FindAllMarkers() function in Seurat and the model-based analysis of single-cell transcriptomics (MAST) algorithm (Finak et al., 2015), as has been applied in previous AD single-cell sequencing studies (Morabito et al., 2020; Yang et al., 2022), unless otherwise specified. We included donor ID as a latent variable to be regressed out in all MAST modeling unless otherwise specified.

#### Endothelial cells vs. non-endothelial cells

We examined gene expression across brain regions in low-pathology endothelial cells using FindMarkers(), opting to retain a gene as an endothelial-specific marker if the average log2 fold change (log2FC) was greater than or equal to 1, if the gene was detected in at least 10% of endothelial nuclei, the p-value was less than 0.05, and the Benjamini-Hochberg (BH)-adjusted p-value was less than 0.1. The resulting set of endothelial marker genes was resupplied to FindMarkers() with no thresholding to calculate the donor-regressed average log2FC across all five cell types for comparison.

#### Region-specific low-pathology endothelial transcriptome analysis

To examine regional endothelial heterogeneity in normal aged brain, we subsetted the integrated endothelial dataset to only the low-pathology subjects (Braak 0/I/II; see Supplementary Table 1) and performed DE analysis using FindMarkers(), with the brain region as the identity variable to compare. Genes were retained as significant for each region if the absolute average log2FC was greater than 0.1, if the gene was detected in at least 10% of endothelial nuclei in the given region, the nominal p-value was less than 0.05, and the BH-adjusted p-value was less than 0.1. These thresholds were chosen based on previously published criteria for endothelial snRNAseq in AD(Lau et al., 2020).

#### Whole-brain high-versus low-pathology endothelial transcriptome analysis

We compared the endothelial transcriptome in the high-versus low-pathology donor cortex, first by combining nuclei from all five brain regions into one analysis. To focus on donors at either end of the neuropathological spectrum, we omitted intermediate pathology donors (B2, n=5) and kept low (B0/1; n=11) and high pathology donors (B3; n=16). DE analysis was performed using FindMarkers() with the pathology level as the identity variable to compare (*i*.*e*. high versus low). Genes were retained as significant if the absolute average log2FC was greater than 0.1, if the gene was detected in at least 10% of endothelial nuclei in the given region, the nominal p-value was less than 0.05, and the BH-adjusted p-value was less than 0.1.

#### Region-specific high-versus low-pathology endothelial transcriptome analysis

Upon identifying genes with differential expression based on pathology level across the whole brain, we further examined genes with region-specific differential expression. We re-ran FindMarkers() for each brain region separately, keeping donor ID as the latent variable to regress out. A gene was retained as significant for a given region if the absolute average log2FC was greater than 0.1, if the gene was detected in at least 10% of endothelial nuclei in the region, the nominal p-value was less than 0.05, and the BH-adjusted p-value was less than 0.1.

#### Identification of gene sets that are temporally related to AD disease stages

To identify genes that were related to the pathological progression of AD, we performed differential expression analysis between adjacent pathology groups (i.e. group 1 vs. group 2; group 2 vs. group 3; group 3 vs. group 4; Supplementary Table 1) using FindMarkers() with region as the latent variable and logistic regression (instead of MAST). Genes with logFC > 0.25 and BH-adjusted p-value < 0.05 and with expression greater than 10% in all groups were included. The standardized gene expression was determined by scaling across all samples for each gene and computing the average score for each pathology group. The standardized gene expression scores were grouped using spectral clustering with SNFTool (v2.3.1; k=6)(Wang et al., 2014).

#### Plaque burden, pTau/Tau and APOE allele endothelial transcriptome analysis

To directly investigate the association between the endothelial transcriptome and AD neuropathology burden, gene expression was modeled as a function of (1) amyloid beta plaque burden, (2) CAA score, (3) pTau/Tau ratio, or (4) APOE E4 allele status, respectively. For all four analyses, data were filtered to only include genes with reads detected in at least 20% of endothelial cells and standardized variance greater than 0.5. For the CAA analysis (model 2), data were additionally filtered to remove samples with missing CAA scores (as indicated by “NA” in the CAA column in Supplementary Table 1). After filtering the data, for each of the four pathology variables, a zero-inflated MAST generalized linear mixed effects model was fit to regress endothelial cell gene expression on the variable of interest using the zlm() function. The donor ID was included as a random effect and the cellular detection rate (“cngeneson”) and percent mitochondrial DNA (“pc_mito”) variables were included as fixed effects, with the method set to “glmer” for a generalized linear mixed-effects model. The ebayes flag was set to FALSE to turn off variance regularization.

### Functional pathway annotation

#### Pathway overrepresentation analysis

To examine which Gene Ontology (GO) biological pathways (BPs) (Carbon and Mungall, 2018) are enriched in endothelial genes of interest, we performed overrepresentation analysis using PANTHER (Mi et al., 2013). The resulting json files were downloaded and imported into R using the jsonlite package (Ooms, 2014). The 36,601 genes assayed in this study were provided as the reference gene set for PANTHER. We defined significance as nominal p-value < 0.05 and adjusted for multiple comparisons in R using the p.adjust() function with BH correction.

#### Gene set enrichment analysis (GSEA)

To complement pathway overrepresentation analysis, we examined the enrichment of gene sets based on the log2FC ranking of all genes using gene set enrichment analysis (GSEA) (Mootha et al., 2003; Subramanian et al., 2005). This method is advantageous in that it does not rely on any specific fold change or significance threshold since it takes the full list of genes (Mootha et al., 2003; Subramanian et al., 2005). A log2FC-ranked list of genes was compiled in the .rnk format and GO BPs were curated in the .gmt format by msigdb (Liberzon et al., 2011). We performed pre-ranked GSEA using the fgseaMultiLevel() function from the fgsea package in R (Korotkevich et al., 2021), with the following parameters: minSize=10, maxSize=500, eps=0.0, nPermSimple=1000. A pathway was considered significantly enriched (positive or negative) if the p-value was less than 0.05 and the BH-adjusted p-value was less than 0.25, as per GSEA recommendations (Subramanian et al., 2005).

## Results

### Identification of endothelial cells from multiple brain regions using single-nucleus RNA sequencing

To examine the region-specific transcriptome of brain endothelial cells in normal control brain and in AD, we performed snRNA-seq on postmortem brain tissue in five regions (EC, ITG, PFC, V2, and V1) available from 16 high-pathology (B3) AD donors, 5 intermediate-pathology (B2) donors, and 11 low-pathology (B0/1) donors (Supplementary Table 1, Figures S1A-B, S2A). We separated out most neurons and oligodendrocytes using fluorescence-based sorting, then clustered nuclei and identified endothelial cell nuclei based on von Willebrand factor (*VWF*) expression, and finally, removed subclusters likely to reflect contamination (see Methods). After these steps, we retained 19,271 genes measured in 51,593 endothelial nuclei (counts per brain region: EC, 3,484; ITG, 9,259; PFC, 10,377; V2, 13,110; and V1, 15,356). An overview of this experimental and analytical workflow is depicted in Figure 1A-B. There was no significant difference in donor postmortem interval (PMI) between B1 versus B3 donors (Figure S1C), and nuclei from the five regions exhibited comparable read counts per nucleus, RNA integrity number (RIN), and total number of genes detected (Figure S2C). Of note, non-endothelial cell types collected are concurrently being analyzed in separate studies(Anon, n.d.; Serrano-Pozo et al., 2022).

**Figure 1.**
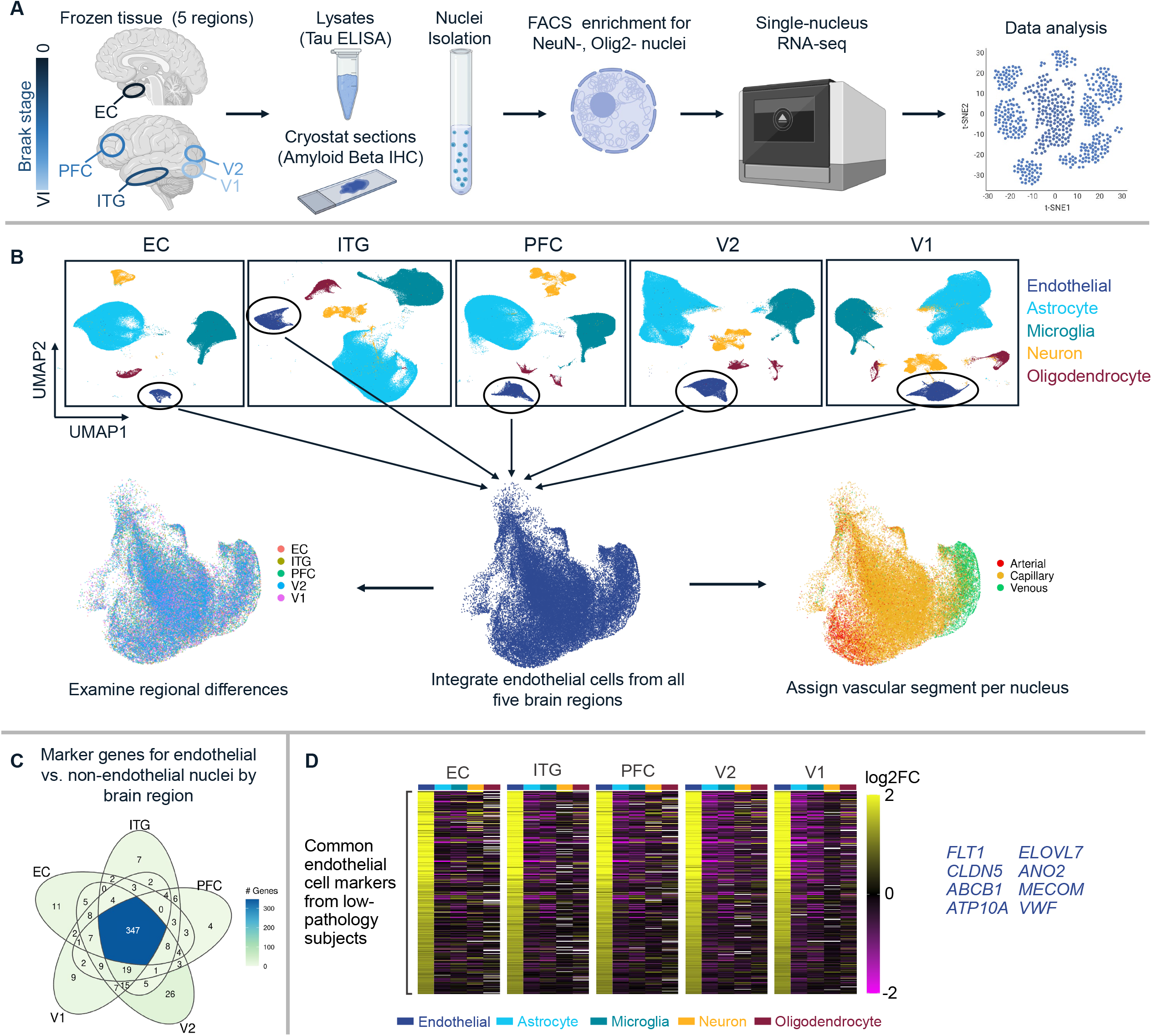
Study design overview. A: NeuN-/Olig2-negative nuclei were isolated from five brain regions (EC, ITG, PFC, V2, V1) and sequenced (10X Genomics) to examine endothelial cell-specific transcriptome throughout the AD brain. B: Endothelial cells were identified in each brain region individually and isolated for integration, enabling the examination of region-specific and vessel segment-specific cerebrovascular transcriptomes. C: Differential expression analysis using Findmarkers() between low-pathology donor endothelial nuclei and low-pathology donor non-endothelial nuclei identified a total of 527 unique endothelial cell markers, 347 of which were shared across all five regions. D: Heatmap visualization of the 347 core endothelial marker genes across cell types depicts the endothelial-specific upregulation. The top eight genes across the regions are highlighted on the right (*FLT1, CLDN5, ABCB1, ATP10A, ELOVL7, ANO2, MECOM*, and *VWF*).

To confirm that the endothelial cell populations were enriched for canonical cell type markers, we compared endothelial nuclei versus non-endothelial nuclei per region (Figure 1C, Supplementary Table 4). Across the five brain regions, a total of 527 unique genes were identified as endothelial cell markers, of which n=347 (65.8%) were shared across all five brain regions. The set of 347 pan-region endothelial marker genes was visualized in a heatmap including other cell types to visually confirm enrichment of endothelial cells in each brain region (Figure 1D, Supplementary Table 5). These include established endothelial cell markers such as *FLT1* (VEGF receptor-1), *CLDN5* (tight junction protein claudin 5), *ABCB1* (ATP-binding cassette sub-family B member 1; P-glycoprotein) and *VWF*. We then characterized the endothelial nuclei by vessel segment zonation, identifying 5,046 arterial, 41,517 capillary and 5,023 venous nuclei, across the five brain regions (Figure S4A). The proportion of capillary nuclei identified here (80.5 percent) is consistent with findings that capillary endothelial cells comprise approximately 85 percent of the blood-brain barrier (Montagne et al., 2017).

### Regional transcriptomic variability in endothelial cells from low-pathology control brains

To investigate regional heterogeneity in the endothelial cell transcriptome, we examined nuclei isolated from aged donors with no to minimal AD-related neuropathology (B0/1 donors in Supplementary Table 1). UMAP visualization of the integrated low-pathology endothelial nuclei at 0.3 resolution did not show any clear differences between regions (Figure 2A). However, in terms of vessel segment composition, there were clear differences in the relative proportions of arterial, capillary, and venous endothelial cells across the five regions (Figure 2B). For example, the EC had more arterial nuclei than the other regions, while V1 had more capillary nuclei than the other regions. Whether this reflects genuine differences in vascular bed composition across the regions or experimental variability remains to be clarified.

**Figure 2.**
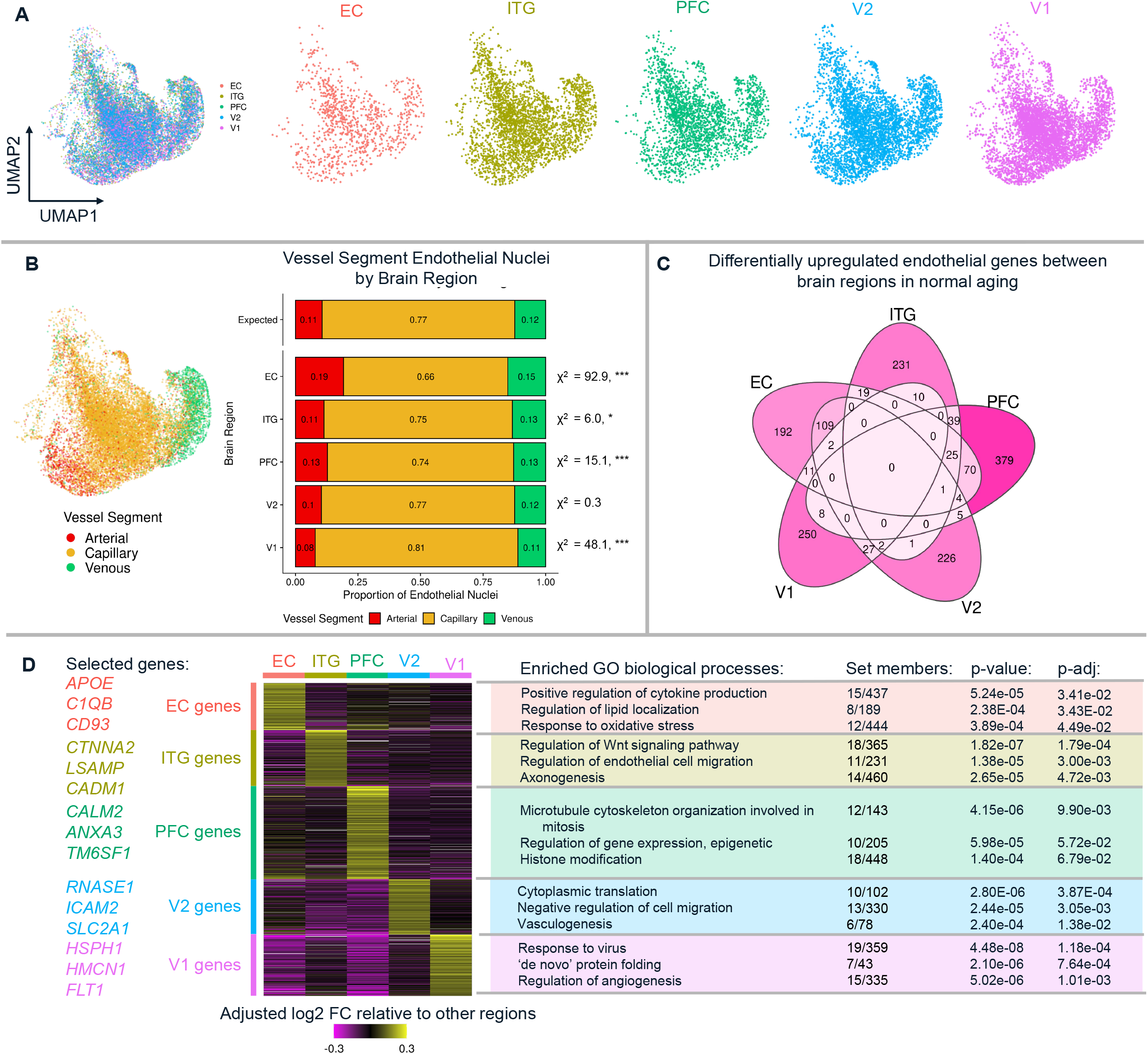
Endothelial cells from low-pathology donor brain regions exhibit differences in transcriptomic pathways and vessel segment composition. A: UMAP visualization of the integrated low-pathology donor nuclei show that the brain regions are evenly distributed in UMAP space. B: UMAP visualization showing endothelial cell arterial-venous zonation of all nuclei. The proportions of nuclei across brain regions differed from expected. Chi-squares test results given on right. * p < 0.05, ** p < 0.005, *** p < 0.001. C: Differential gene expression analysis showing upregulated genes in each of the five brain regions. D: Heatmap showing upregulated genes for each brain region are increased relative to other areas and Gene Ontology (GO) biological process (BP) overrepresentation analysis identified the pathways that are enriched in each region’s marker gene sets

We then identified whether individual genes varied across brain regions and identified between 192 and 379 genes (or 1-2% of all genes detected in endothelial cells) that were uniquely upregulated in one of the regions relative to the other four, making them region-specific marker genes (Figure 2C). Critically, donor ID was included as a latent variable in this, and all subsequent, analyses. Relevant region-specific genes included *APOE, C1QB* and *CD93* in EC; *CTNAA2, LSAMP*, and *CADM1* in ITG; *CALM2, ANXA3*, and *TM6SF1* in PFC; *RNASE1, ICAM2*, and *SLC2A1* in V2; and *HSPH1, HMCN1*, and *FLT1* in V1. These regional marker genes were then grouped into biological pathways (BPs)(Figure 2D, Supplementary Table 6,7). EC marker genes were enriched for cytokine production, regulation of lipid localization and oxidative stress response. ITG marker genes were enriched for Wnt signaling, endothelial cell migration, and axonogenesis. PFC marker genes were enriched for microtubule organization, epigenetic gene expression, and histone modification. V2 marker genes were enriched for cytoplasmic translation, downregulation of cell migration, and vasculogenesis. V1 marker genes were enriched for virus response, “de novo” protein folding, and angiogenesis regulation. These data together suggest that differences in baseline gene expression exist in each of the five regions examined in the aging brain even before AD pathology develops.

### AD is associated with unique sets of dysregulated gene networks in endothelial cells

To examine AD related changes in each brain region, we first analyzed AD-associated transcriptomic differences in endothelial cells from all five brain regions combined by comparing high-pathology (B3) vs. low-pathology (B0/I) donors. We identified 936 genes that were upregulated in high-pathology donor endothelial nuclei and 962 that were downregulated (Figure 3A, Supplementary Table 8). Among the top ten upregulated genes, eight of them are heat shock family proteins (HSPs) with log2 fold-change (log2FC) values of 2.18 – 1.04, representing more than double the expression of these genes in high pathology AD individuals compared to low-pathology controls. Less extensive fold-changes were observed among the top ten downregulated genes, which had log2FC values of -0.8 – -0.51. To validate the results of this analysis, we performed immunohistochemistry in sections from the temporal and visual cortices of donors used in this study. To validate expression changes, we selected AD-upregulated genes *MT2A, CREM, IGFBP3, DST* and *HSP90AA1*, which represent a range of small to large effect sizes, and confirmed expression at the protein level in capillary endothelial cells(Figure 3B). No specific cortical layer differences were observed in the expression of these proteins by histology.

**Figure 3.**
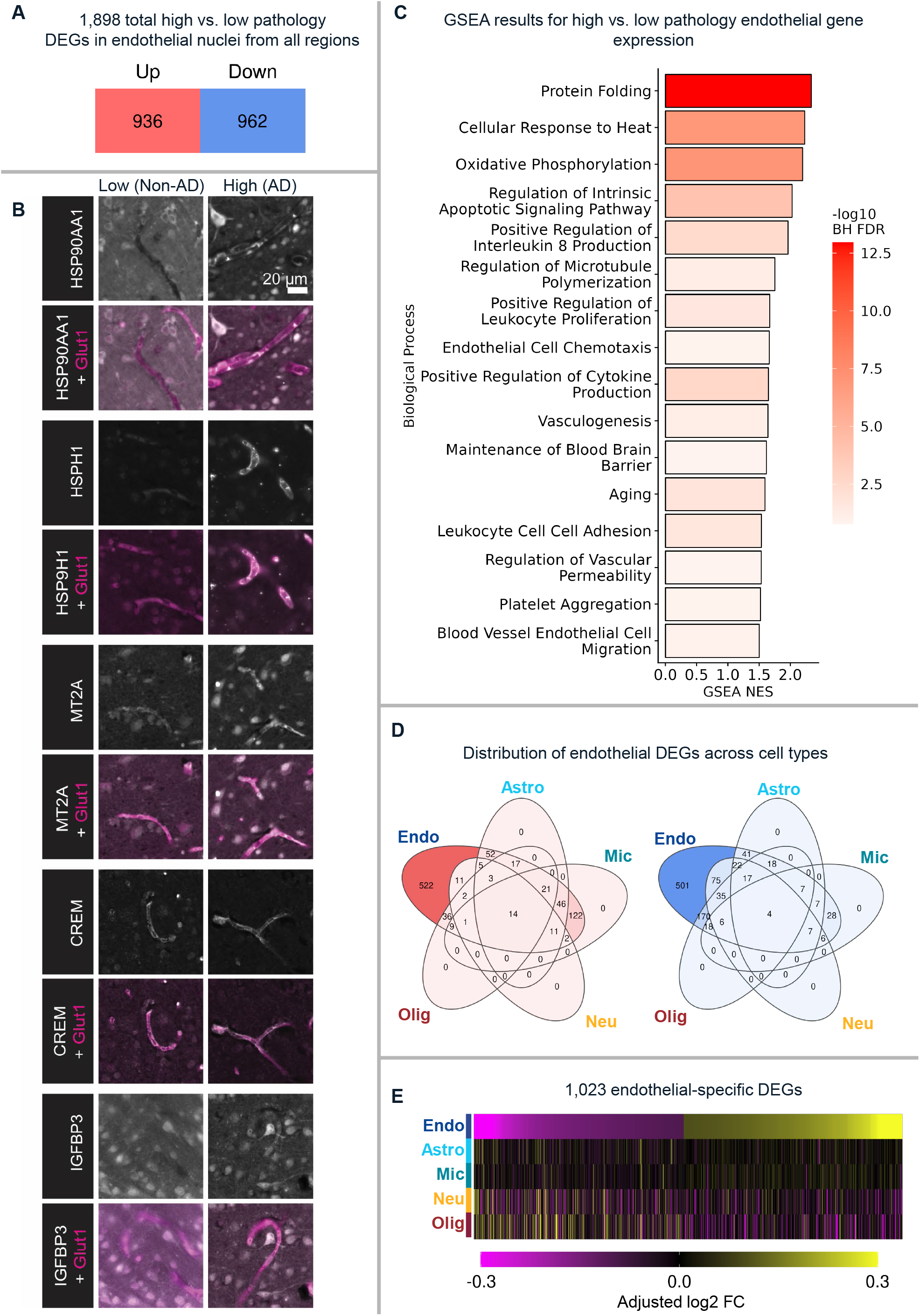
Whole-brain differential expression analysis reveals dysregulated pathways that are specific to endothelial cells in the AD brain. A: In high versus low pathology endothelial nuclei, there are 936 upregulated genes and 962 downregulated genes. B: Immunofluorescence images of five representative genes that are upregulated in AD brain endothelial cells (white). Glut1 co-labeling is used to identify endothelial cells (magenta). All images are from V1/V2 with the exception of MT2A which is from EC. C: GSEA reveals that high-pathology endothelial cells are enriched for several pathways including those related to protein folding, IL-8 production, leukocyte adhesion, and blood-brain barrier maintenance. Only a subset of GO BPs are shown in this figure. NES=normalized enrichment score. D: Differential expression analysis in each cell type shows that 522/936 of the upregulated genes and 501/962 of the downregulated genes are specific to endothelial cells. E: Heatmap visualization of these 1,023 genes depicts their specificity to endothelial cells in AD.

Next, we examined whether endothelial cells from high-pathology donors were enriched for different BPs compared to low-pathology donors using GSEA. For each GO BP, we calculated its normalized enrichment score (NES), which indicates how much the genes in a given gene set are over-represented at the top or bottom of the entire ranked list of genes, normalized to gene set size. Among all 19,271 genes, a total of 660 BPs were significantly positively enriched in the high-pathology endothelial cells, including upregulated pathways related to protein folding, cytokine production, BBB maintenance, and leukocyte adhesion (Figure 3C, Supplementary Table 9). Of the 660 BPs, 68 pathways were downregulated which included those related to lipid and glycoprotein metabolism.

Previous studies that examined the AD cerebrovascular transcriptome have not clarified the endothelial cell specificity of resulting gene expression changes. To address this, we calculated the average high-versus low-pathology log2FC for the n=1,898 genes from Figure 3A for each of the other four cell types (astrocytes, microglia, neurons, and oligodendrocytes). This revealed that 522 of the upregulated genes and 501 of the downregulated genes are unique to endothelial cells (Figure 3D-E, Supplementary Table 10). Of the genes that were not unique to endothelial cells, most of the upregulated genes were shared with microglia (n=122) and most of the downregulated genes with oligodendrocytes (n=170). Of note, 32 of the upregulated genes unique to endothelial cells were identified in Uniprot to be secreted proteins and may thus represent potential plasma or cerebrospinal fluid (CSF) biomarkers for brain endothelial cell changes. These include *ADM, BMP6, EGFR, IL32, IGFBP3, TGFB1*, and *WNT2B*.

### Brain regions exhibit different endothelial transcriptomes amid AD pathology

Since the five brain regions assayed are at different stages along the Braak continuum(Braak and Braak, 1991b; Braak et al., 2006b), we next asked whether BPs that were identified from analysis of total endothelial nuclei were more specific to individual regions. We re-examined these 16 pathways for each region (shown in Figure 3C), comparing their NES for high pathology versus low pathology subjects (Figure 4A, Supplementary Table 11). Of the sixteen BPs, seven were significantly enriched in all five regions: protein folding, oxidative phosphorylation, cellular response to heat, regulation of intrinsic apoptotic signaling pathway, vasculogenesis, aging and positive regulation of cytokine production. Of note, maintenance of the blood brain barrier was significantly enriched in PFC, ITG and to a lesser extent in V1, but not in V2 or EC.

**Figure 4.**
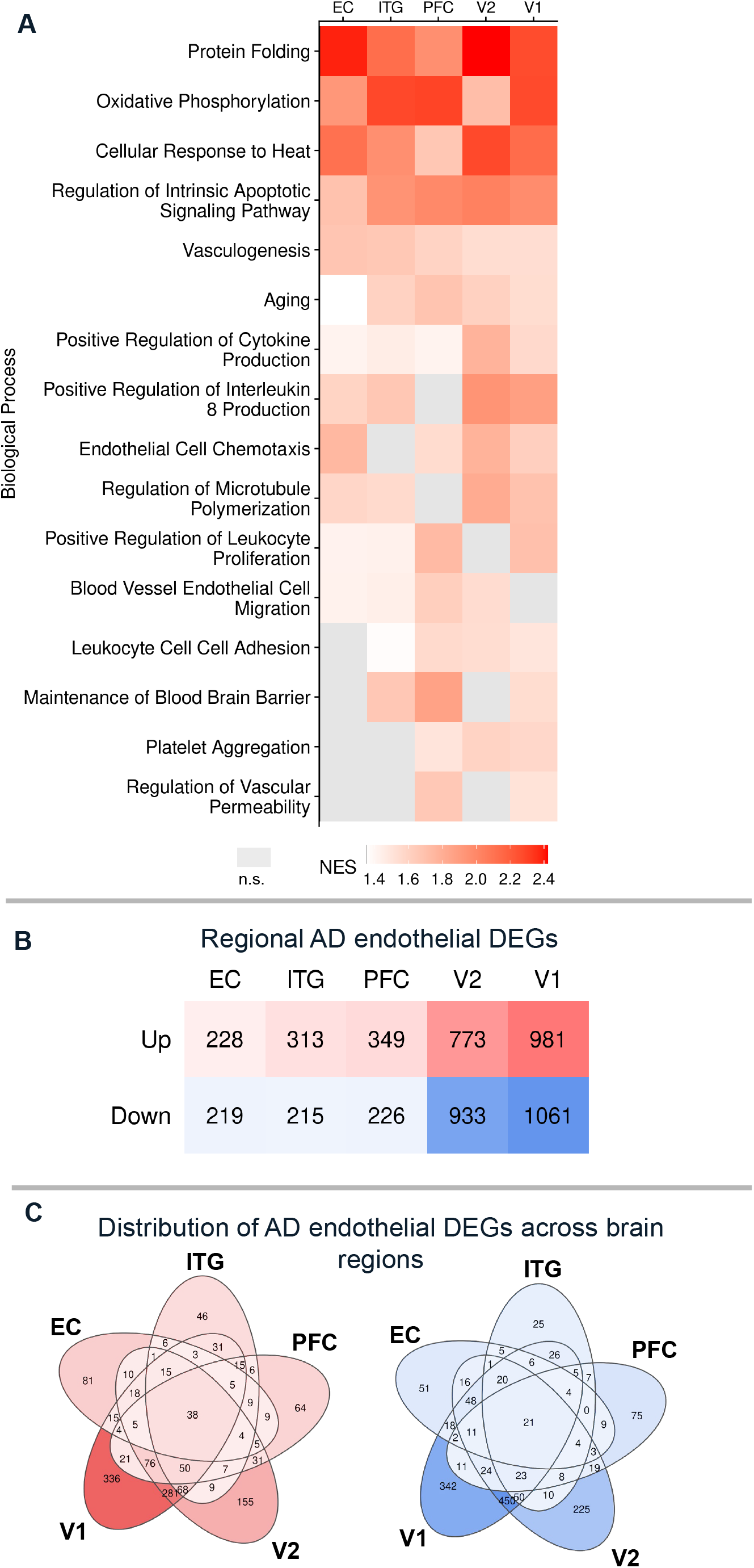
Regional differences in endothelial cell responses to AD pathology. A: The same pathways highlighted in Figure 3C are shown in region-wise GSEA comparing high versus low pathology donors, showing that some pathways are enriched in all five regions (*e*.*g*. protein folding, vasculogenesis) while others are more region-specific (*e*.*g*. blood brain barrier maintenance, regulation of vascular permeability). NES=normalized enrichment score. B: The number of differentially expressed genes increases from EC to V1 along the Braak stage continuum. C: Euler diagram visualization of the genes of interest across regions shows that most genes are uniquely dysregulated in one or two regions, while a few are commonly dysregulated in all five regions (38 upregulated and 21 downregulated).

We further examined the AD-related endothelial transcriptome within each region separately. Region-wise gene expression changes appeared to increase along the Braak stage neuroanatomical continuum, with the lowest number in the EC (228 up, 219 down) and the highest number in the V1 (981 up, 1061 down) (Figure 4B, Supplementary Table 12). Nearly all genes were protein coding genes; of these, 13-33% were exclusive to one region, while others were shared between many or all regions (Figure 4C). These included the upregulation of 38 genes enriched for detoxification and metal ion response (metallothionein genes *MT1E, MT1F, MT1M, MT1X and MT2A*) and notably included *PICALM*, which has been previously associated with AD risk in genome-wide association studies (GWAS)(Harold et al., 2009). The 21 common downregulated genes included ones related to inhibition of TGF-beta signaling (*SMAD6, SMAD7*), endocytosis (*RFTN1, MYOF*) and enzymes that modify basement membrane and glycocalyx (*HGSNAT, CHST15, SULF2, GALNT17*). These AD-related genes showed limited overlap with the region-specific genes (i.e., a gene identified in low pathology nuclei as a marker of V1 endothelial cells was not also altered throughout the brain with AD pathology). Taken together, these findings indicate that endothelial cells within specific brain regions express diverse biological pathways in AD.

### Endothelial cell gene expression programs exhibit diverse temporal relationships with disease progression

One of the limitations of human tissue studies is the difficulty in resolving the temporal relationships between disease stage and transcriptomic changes. This study was designed *a priori* to help address this by including donor samples that fall into four equally sized (n=8) groups based on Braak stage (0-VI) and CERAD score (neuritic plaques; Supplementary Table 1). Pair-wise comparisons between adjacent groups yielded genes that were increased or decreased with pathological burden, and clustering of these genes revealed six common trends in gene expression with disease progression (Figure 5, Supplementary Table 14). Some of these genes showed trajectories that decreased linearly with pathology including ion transport and intracellular signal transduction (Gene Set #1), while others were enriched for pathways that increased linearly including genes related to metal ion homeostasis, apoptosis, and cell cycle regulation (Gene Set #2). By comparison, genes related to lipid localization and cytoskeleton organization appeared relatively consistent across disease stages (Gene Set #3). Gene Set #4, which includes protein folding, cellular response to heat, and regulation of proteolysis genes also showed increases in intermediate stage pathology donors but then returned to baseline in late-stage disease. Other transient changes in intermediate states were observed in Gene Set #5, which included antigen processing, calcium ion response, and vasculature development genes. Lastly, Gene Set #6 included vascular transport and cell signaling genes that were only reduced in an intermediate stage. In all, these data indicate that endothelial cell gene expression changes follow complex, non-linear trajectories in response to disease progression.

**Figure 5.**
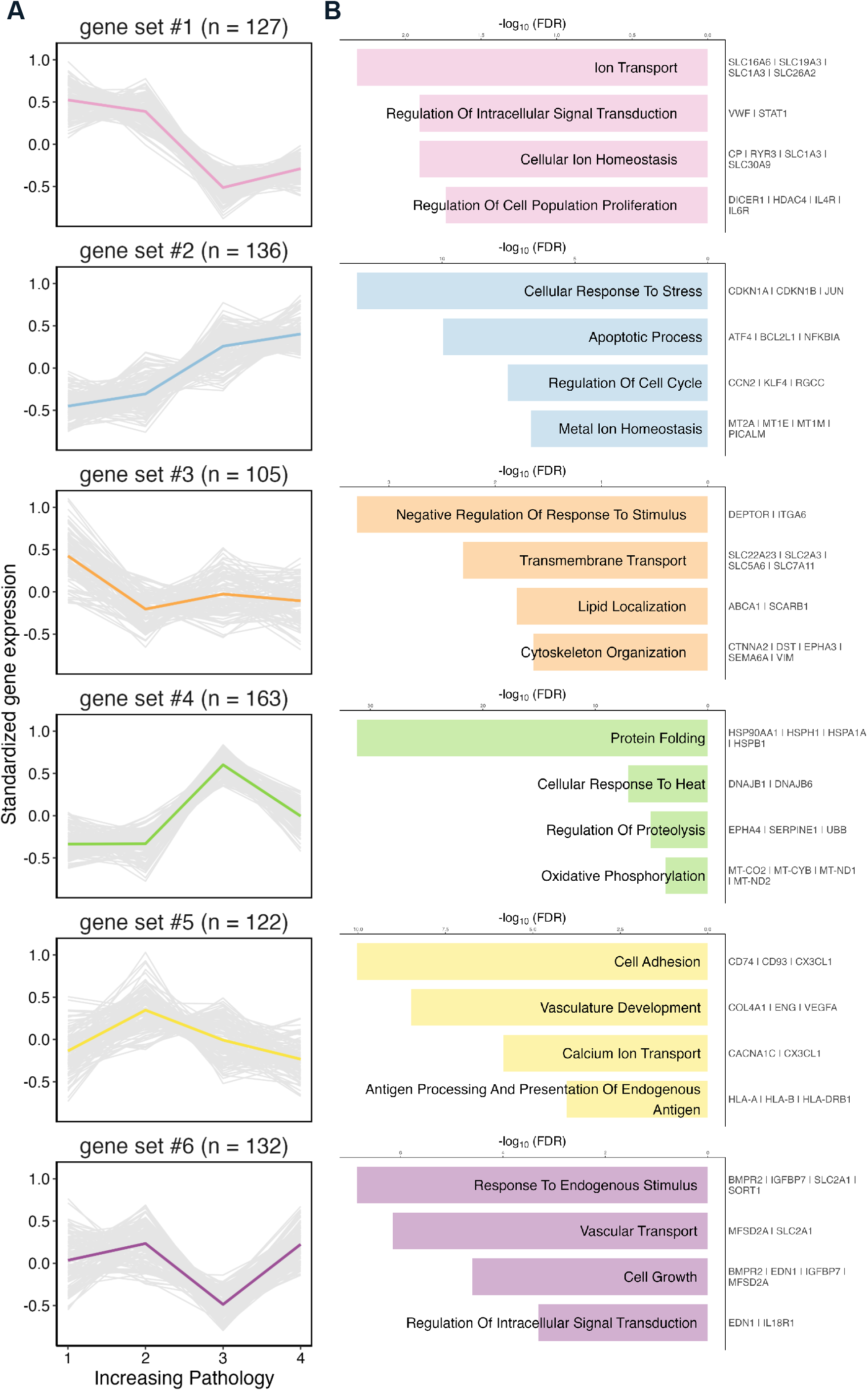
Temporal differences in endothelial cell responses to AD pathology. **(A)** Six gene sets identified by pair-comparisons between tissue from four distinct donor categories (1=Braak 0-II, no neuritic plaques; 2=Braak II/III, sparse-moderate neuritic plaques; 3=Braak V, moderate neuritic plaques; 4=Braak VI, severe neuritic plaques). The graphs show standardized gene expression on the y-axis across each of the four disease stage categories. Individual genes are plotted as grey lines and the colored line indicates the mean trend for each gene set. (B) Selected top GO biological process terms plotted on the left with example genes from each set listed on the right.

### Distinct endothelial gene changes are related to amyloid plaque burden, CAA, tau and APOE genotype

To examine how these endothelial gene expression changes relate to AD neuropathology, we performed linear regression analysis of individual gene expression on histological amyloid plaque burden, phosphorylated tau content, and *APOE* genotype. With regards to amyloid beta plaque load (measured by 3D6-positive area fraction via immunohistochemistry on adjacent tissue sections), 861 genes were found to be positively or negatively associated with plaque burden including 89 that were significant after correcting for multiple comparisons (Figure S5A). In this subset, top genes that were positively associated with amyloid plaque burden include *TMSB10, EEF1A1, TPT1, KLF2, NFKBIA*, and *ICAM2*, and genes that were negatively associated with amyloid plaque burden include *CTNNA2, GLIS2, FBXL7, PDZRN3, FHIT*, and *ABCA1* (Supplementary Table 15). BPs that were overrepresented in this gene set were related to metabolism, inhibition of apoptosis, zinc homeostasis, translation, and endosomal transport (Supplementary Table 16).

We also examined whether endothelial gene expression was affected by cerebral amyloid angiopathy (CAA; scored 0 to 4 based on severity using paraffin embedded sections from the opposite hemisphere). This yielded 684 genes that were positively or negatively associated with CAA, including 37 that remained significant after corrections for multiple comparisons (Figure S5B, Supplementary Table 17). This included positively associated genes *BACE2* and *LRP1B* and negatively associated genes *SLC7A5, SLC16A1, SLC2A1, TFRC, FTH1 and CLU*. Of the CAA-related genes with nominal p-values <0.05, only 11 positively associated genes and 34 negatively associated genes overlapped with those related to amyloid beta plaques. CAA-related genes included overrepresented BPs for regulation of amyloid beta formation, transport across the blood-brain barrier, iron homeostasis and response to VEGF stimulus (Supplementary Table 18). Collectively, this suggests that CAA-related changes are distinct from changes related to amyloid beta plaques.

Finally, we examined the relationship between gene expression and phosphorylated tau (measured biochemically as a ratio between p231 tau and total tau in the same bulk tissue used for the snRNA-Seq) or *APOE* genotype (E3 *vs*. E4). This resulted in 154 and 177 genes associated with these variables respectively, but none were statistically significant after correction for multiple comparisons (Supplementary Tables 19, 20). Further, no BPs were overrepresented in these gene sets. Altogether, few of the genes associated with amyloid plaque burden, CAA, phosphorylated tau, and *APOE* overlapped, and no genes correlated with all four variables. This indicates that AD-related neuropathological factors have distinct effects, and that amyloid beta exerts the greatest impact on endothelial cell gene expression independent of brain region.

### High-tau endothelial cells recapitulate published enrichment of senescence, endothelial aging, and CAA pathways

To anchor the findings presented herein with previous studies examining the cerebral vasculature in health and disease, we curated set of genes identified in stroke, CAA, aging, and small vessel disease (SVD) and compared them to this dataset (Supplementary Table 21). At the whole-brain level, high-vs. low-pathology endothelial cells were significantly enriched for genes related to CAA (Moursel et al., 2018), endothelial aging (Zhao et al., 2020), senescence response (Dehkordi et al., 2021), and the endothelial senescence-associated secretory phenotype (SASP) (Kiss et al., 2020) (Figure 6A-B). These cells also showed a trend (nominal p-value < 0.05, adjusted p-value < 0.1) for enrichment of genes involved in senescence initiation (Dehkordi et al., 2021). At the regional level, two pathways remained significantly enriched across all five regions: CAA and endothelial aging (Figure 6C) while senescence-related pathways were more region-specific, particularly in the ITG and PFC. Genes associated with stroke, white matter hyperintensities, small vessel disease and other disorders (progressive supranuclear palsy and frontotemporal dementia) were not enriched. Within these donors, these data confirm key endothelial changes observed in previous datasets and highlight the importance of considering regional differences when examining specific pathways.

**Figure 6.**
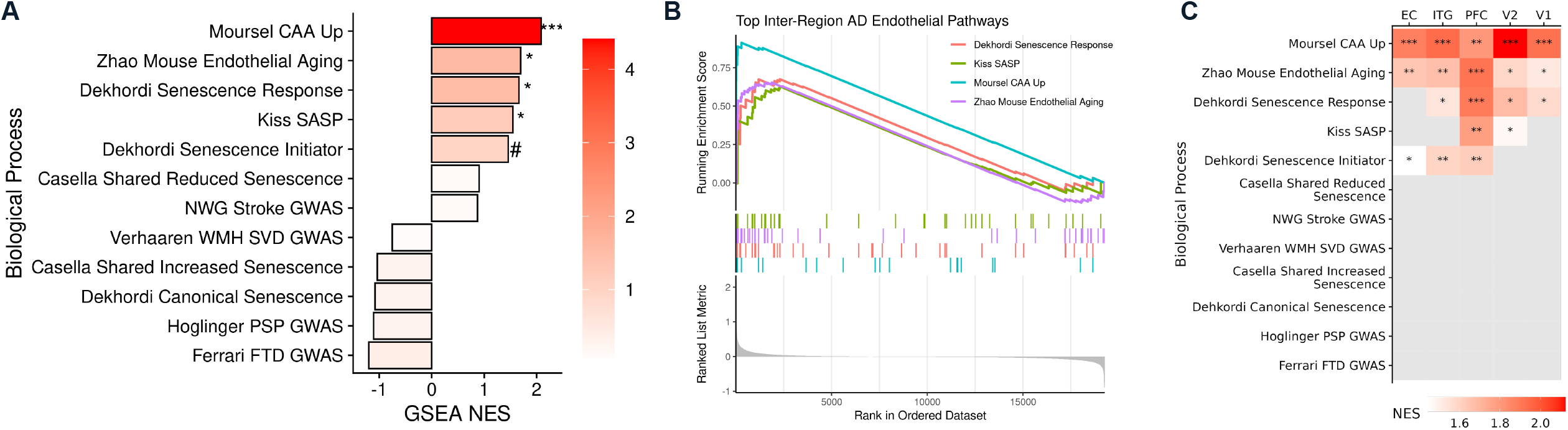
Enrichment of aging and disease-associated pathways in endothelial cells. Comparison with previous publications highlights that high-pathology donor endothelial cell nuclei are enriched for gene sets related to cerebral amyloid angiopathy (CAA), endothelial aging, senescence response, and the senescence-associated secretory phenotype (SASP). A: Normalized enrichment scores (NES) are shown for each of the twelve gene sets derived from recent AD and/or vascular disease-related publications. All brain regions were included in this analysis. B: GSEA plot visualization depicts the genes’ rank in the ordered dataset versus running enrichment score in each of the significantly enriched modules, respectively. C: Region-wise GSEA reveals that CAA and endothelial aging gene sets are enriched in endothelial cells from all five regions, while senescence response, SASP, and senescence initiator pathways are region-specific. * p < 0.05, ** p < 0.005, *** p < 0.001, # p < 0.05 but BH-adjusted p > 0.25. NES=normalized enrichment score. Running enrichment score plots are shown in supplementary Figure S6.

## Discussion

Using a large dataset of endothelial cell nuclei, the transcriptomic analysis that we report reveals new insights into regional gene expression differences in normal aged brain and AD. These data indicate that endothelial cells from different cortical areas are clearly distinguishable and are characterized by the expression of different marker genes and biological pathways. We also observed that AD pathology affects gene expression in brain endothelial cells, particularly those that make up capillaries. By comparing across donor samples from multiple disease stages, we observed the temporal relationship between pathology and endothelial transcriptome changes. Further, using this data we compared the extent that AD-related factors, including plaques, CAA, tau and APOE genotype, impact this cell type and examined how previously reported transcriptomic signatures vary between brain regions.

A fundamental biological question we set out to answer was whether the vascular transcriptome differs across brain regions. Previous endothelial cell transcriptomic datasets have been derived from mouse models (He et al., 2018; Vanlandewijck et al., 2018), where examination of distinct cortical areas is challenging, and existing human data sets include only a single cortical area per donor (Lau et al., 2020; Yang et al., 2022). By incorporating multiple cortical areas per donor, we showed that significant heterogeneity exists in endothelial cells, with 1-2% of total genes detected in endothelial cells differentially expressed in each region. Previous assays of transcriptomic differences that used mouse endothelial cells from brain, heart, and lung showed that differential expression of 3-9% of the total genes was responsible for tissue specificity (Jambusaria et al., 2020). Similarly, mouse and human brain endothelial cells show a 4.4% divergence in total gene expression (Yang et al., 2022). This suggests that the number of genes with expression differences across brain regions in the normal aged brain could have a relatively large impact on endothelial cell phenotypes. This warrants further study to better understand functional implications of this regional heterogeneity in the healthy endothelial transcriptome.

A better understanding of these transcriptomic changes could help explain why certain brain areas are more prone to CAA or neurodegeneration. Interestingly, visual cortex areas, which are affected late in AD progression and experience less neurodegeneration, expressed more genes related to vasculogenesis and angiogenesis. This transcriptomic difference might explain the relatively greater vascularization and distinct organization of visual cortex versus other brain areas (Duvernoy et al., 1981; Schmid et al., 2019). By comparison, highly vulnerable areas such as the entorhinal cortex expressed more oxidative stress-related genes in normal aged brain, suggesting endothelial dysfunction in this region even in the absence of severe AD pathology. In sum, these data support the idea that important differences in endothelial beds exist among brain regions. Whether these are due to development (*i*.*e*. origination of capillaries from the posterior cerebral artery versus middle cerebral artery), local structure or age-related changes in various brain areas remains to be examined.

In comparing donor tissue from high- and low-pathology brains, the most highly upregulated genes were members of heat shock family (HSP) proteins. HSPs are important components of translational machinery in endothelial cells, and such upregulation could be a response to stress stimuli including inflammatory cytokines, oxidized LDL, elevated blood glucose, and fluid sheer stress(Hochleitner et al., 2000). Exposure of cultured endothelial cells to metal ions is also known to result in upregulation of HSPs (Wagner et al., 1999). Increased expression of these proteins could also indicate a switch from quiescence to an activated, proliferative state (Pawlowska et al., 2005). Importantly, these genes appeared to have distinct trajectories and increased in intermediate disease stages only to return to baseline in late-stage disease. Because HSPs are a critical component of proteostasis mechanisms, we speculate that their elevation throughout the AD brain raises the possibility of brain-wide proteostatic stress that impacts cells without visible protein aggregates and that failure of this pathway in late-stage disease may contribute to severe phenotypes. Considering that these genes were increased in all brain areas examined, and not significantly associated with amyloid plaque burden, CAA, phosphorylated tau levels, or *APOE* genotype, it is possible that this is a response to neurodegeneration more generally. Future studies should expand the scope to non-AD neurodegenerative diseases like frontotemporal dementia or Lewy body disease to clarify the specificity of AD in dysregulated expression of these pathways.

We also explored regional heterogeneity in response to AD pathology and observed that endothelial responses to pathology are not uniform across brain regions, with a greater number of gene expression differences in the visual cortex compared to other areas. One explanation could be more age- and/or disease-related pathology accumulation in other brain areas like the EC, resulting in more similarities and fewer gene expression differences between normal aged brain and AD. It is notable that the Moursel *et al*. CAA gene set was prominently enriched in AD endothelial cells, and that CAA is more frequently encountered in occipital regions such as the visual cortex compared to anterior areas (Vinters and Gilbert, 1983; Moursel et al., 2018). This is also in line with our finding that both CAA and amyloid plaque burden are more strongly associated with endothelial gene expression changes than tau pathology and/or the presence of the *APOE4* allele in this dataset. However, future studies that include greater numbers of *APOE4* carriers, including AD *APOE4* homozygotes and control subjects without AD, will be required to specifically address the contribution of this risk allele to endothelial cell transcriptomic changes.

Lastly, it was surprising that genes associated with amyloid beta plaques minimally overlapped with genes associated with CAA in this dataset. Both CAA and amyloid beta plaques frequently, but not always, co-occur within regions (Vinters et al., 1996; Thal et al., 2003). This indicates that while both result in local increases in pathological amyloid beta, they appear to exert different effects on endothelial transcription programs. The form of pathological amyloid beta, vessel segment, and specific location within cortex may contribute to this observation. We note that CAA predominantly affects leptomeningeal vessels (excluded from this analysis via gross dissection) particularly near the occipital cortex, while plaques are often found more diffusely in deeper cortical layers (Rogers and Morrison, 1985; Charidimou et al., 2017). As such, future studies that examine both pial vessels and parenchyma using spatial transcriptomics could determine how local microenvironments contribute to the observed gene expression programs in endothelial cells.

The current study extends and reinforces our earlier work examining gene expression changes in isolated vasculature, and work from other groups that have performed single-cell analysis using similar preparations (Lau et al., 2020; Yang et al., 2022). Previously, we had observed that with increasing severity of AD pathology, isolated blood vessels expressed more cellular senescence-related changes as measured by qPCR (Bryant et al., 2020). The present work shows that senescence-related gene signatures are increased across several brain regions and confirms these changes in endothelial cells in the absence of other vascular cell types. Senescent cells are known to have paracrine effects on surrounding tissue, often inducing sprouting of new dysfunctional blood vessels. Consistent with this, other changes in the proliferative capacity of endothelial cells were also noted, including the enrichment of genes related to vasculogenesis at the whole-brain level. Previous reports have observed altered angiogenesis, BBB maintenance, and leukocyte adhesion gene changes in samples from prefrontal cortex (Lau et al., 2020). Our analyses add to this and show that vasculogenesis- and angiogenesis-related genes have distinct regional and temporal expression profiles.

Altogether, these data can be used to guide future studies exploring disease processes in endothelial cells and indicate that careful tissue selection is warranted, as cortical regions are not all equivalent and show key differences at baseline in the aged brain and in response to AD pathology. We found that amyloid beta plaques and CAA have the greatest impact on gene expression programs in endothelial cells from AD donors. Lastly, while endothelial cells are not typically associated with protein aggregation, upregulated protein folding pathways suggest that proteostatic stress is a key pathway in this cell type.

## Acknowledgements

This work was supported by NIH grants R01AG071567 (Bennett), P30AG062421 (Hyman and Das), K08AG064039 (Serrano-Pozo) and MASSCATS. We would like to acknowledge individuals who have assisted in completing this project including Patrick Dooley and Teresa Connors for their help with tissue selection, Derek Oakley and Matthew Frosch for their neuropathological expertise, and Simon Dujardin and Sarah Hopp for their intellectual support. We also thank the many donors and their families who have contributed to the Massachusetts Alzheimer’s Disease Research Center.

## Data availability

All raw sequencing data is deposited and available in the NCBI Gene Expression Omnibus (GEO) under accession code to be provided at publication. Data may also be explored via an interactive web browser at: http://alzdatalens.partners.org/ (Datalens).

## Supplemental Figures

**Figure S1**. Neuropathological features are shown for each donor and brain region. A: The region-wise pTau:Tau ratio is plotted for each donor, labeled by ABC-B. B: The region-wise amyloid plaque area is plotted for each donor, labeled by ABC-C. Donors are ordered from greatest to least severity (left to right) in each panel. C: Postmortem interval (PMI) for donors by ABC-B pathology group. Values were compared using the Kruskal-Wallis nonparametric analysis of variance (p=0.57).

**Figure S2**. QC for endothelial cells sequenced. A: Summary of the number of donors with tissue for each brain region, stratified by ABC-B group. B: *VWF* expression overlaid onto the UMAP visualization of all nuclei highlights the endothelial nuclei cluster for each brain region. C: Standard quality control metrics plotted for each brain region confirm comparable QC across regions. D: Comparison of the total number of endothelial nuclei per donor (top row) and the proportion of endothelial cells to total nuclei isolated per donor (bottom row) across the Braak stages per brain regions. Values were compared using the Kruskal-Wallis nonparametric analysis of variance, with n.s. indicating p > 0.05.

**Figure S3**. QC for Seurat integration. A: Elbow plots for the five brain regions show the standard deviation explained by each of the first 50 principal components (PCs). B: Cell type marker visualization for the subclusters identified in endothelial cells integrated from all five regions identifies subclusters with heightened non-endothelial contamination (4, 5, 7, 10). Ast = astrocyte; mic = microglia; oligo = oligodendrocyte; end = endothelial cell; opc = oligodendrocyte precursor cell; per = pericyte.

**Figure S4**. Vessel zonation QC analysis. A: The number of nuclei and median number of genes detected in each vessel segment are shown. B: Standard quality control metrics are plotted for each vessel segment, confirming comparable sequencing quality across segments. C: Differential expression analysis in high-versus low-pathology endothelial nuclei stratified by vessel segment reveals that the vast majority of genes of interest arise from capillary nuclei.

**Figure S5**. Genes associated with amyloid beta plaques and CAA. A: Examples of low and high amyloid plaque burden across brain regions and volcano plots showing genes that are positively or negatively associated plaque burden. B: Examples of CAA grading severity (0-3) across 4 donors and volcano plots showing genes that are positively or negatively associated CAA severity.

### Supplemental Tables

**Supplementary Table 1**. Clinical and neuropathological characteristics of the n=32 donors included in this study. In the five rightmost columns, an X indicates that nuclei were sequenced for the given donor and region, while a blank space means tissue was unavailable for sequencing.

**Supplementary Table 2**. Marker genes used to identify cell types.

**Supplementary Table 3**. Markers used to identify vessel segments. Gene list adapted from Yang et al. 2022.

**Supplementary Table 4**. Genes significantly upregulated in endothelial cell nuclei relative to other cell types. (Data shown in Figure 1C)

**Supplementary Table 5**. Comparison of endothelial marker genes expression in other cell types (Data shown in Figure 1D).

**Supplementary Table 6**. Differentially upregulated endothelial cell markers between brain regions in low-pathology subjects. (Data shown in Figure 2D).

**Supplementary Table 7**. Region-specific GO Biological Processes (BPs) identified from GSEA on expression levels from Table S6.

**Supplementary Table 8**. Differential expression analysis for high-versus low-pathology endothelial cells, all brain regions together.

**Supplementary Table 9**. Pathology-associated GO BPs identified from GSEA on expression levels in Table S8.

**Supplementary Table 10**. Expression levels across cell types for genes uniquely up-or down-regulated in high-pathology endothelial cells

**Supplementary Table 11**. Pathology-associated endothelial-specific GO BPs identified from GSEA on expression levels in Table S10.

**Supplementary Table 12**. Differential expression analysis for high-versus low-pathology endothelial cells, separated by brain region

**Supplementary Table 13**. Pathology-associated endothelial- and region-specific GO BPs identified from GSEA on expression levels in Table S12. (Data shown in Figure 4A).

**Supplementary Table 14**. Gene sets associated with AD pathology temporal dynamics.

**Supplementary Table 15**. Genes associated with amyloid plaque burden.

**Supplementary Table 16**. Table containing amyloid plaque-associated GO BPs obtained from enrichment analysis.

**Supplementary Table 17**. Genes associated with CAA pathological burden.

**Supplementary Table 18**. Table containing CAA-associated GO BPs obtained from enrichment analysis.

**Supplementary Table 19**. Genes associated with ptau/tau levels.

**Supplementary Table 20**. Genes associated with presence of the APOE4 allele.

**Supplementary Table 21**. List of gene sets related to aging and neurodegenerative disease that were compared to this dataset.

