## Supplemental Figures for "Endothelial Cells are Heterogeneous in Different Brain Regions and are Dramatically Altered in Alzheimer’s Disease"

Figure S1

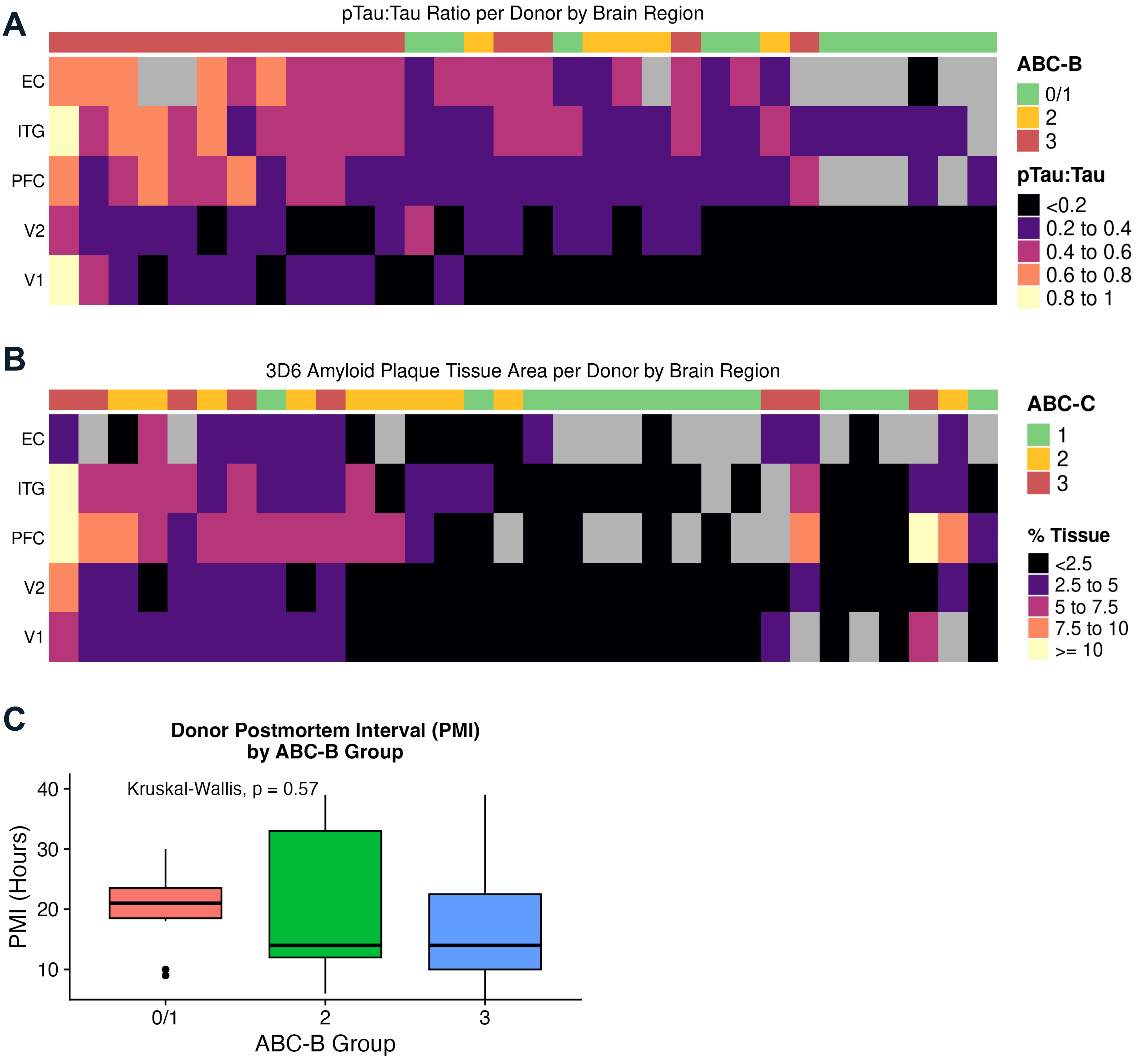
A
pTau:Tau Ratio per Donor by Brain Region
EC
ITG
PFC
V2
V1
ABC-B
0/1
2
3
pTau:Tau
<0.2
0.2 to 0.4
0.4 to 0.6
0.6 to 0.8
0.8 to 1
B
3D6 Amyloid Plaque Tissue Area per Donor by Brain Region
EC
ITG
PFC
V2
V1
ABC-C
1
2
3
% Tissue
<2.5
2.5 to 5
5 to 7.5
7.5 to 10
>= 10
C
Donor Postmortem Interval (PMI)
by ABC-B Group
Kruskal-Wallis, p = 0.57
PMI (Hours)
40
30
20
10
0/1
2
3
ABC-B Group

Figure S2

**A**

| Region | B1 | B2 | B3 |
|---|---|---|---|
| EC | 6 | 4 | 12 |
| ITG | 10 | 5 | 16 |
| PFC | 7 | 5 | 16 |
| V2 | 11 | 5 | 16 |
| V1 | 11 | 4 | 16 |
| **TOTAL** | **11** | **5** | **16** |

**B**

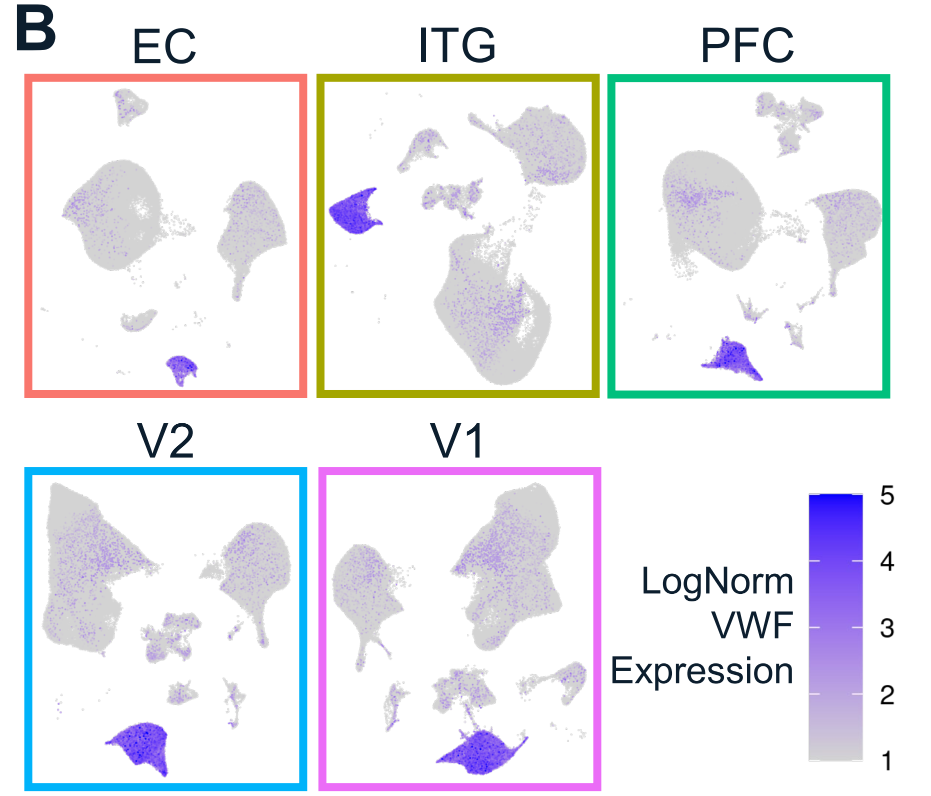


**C**

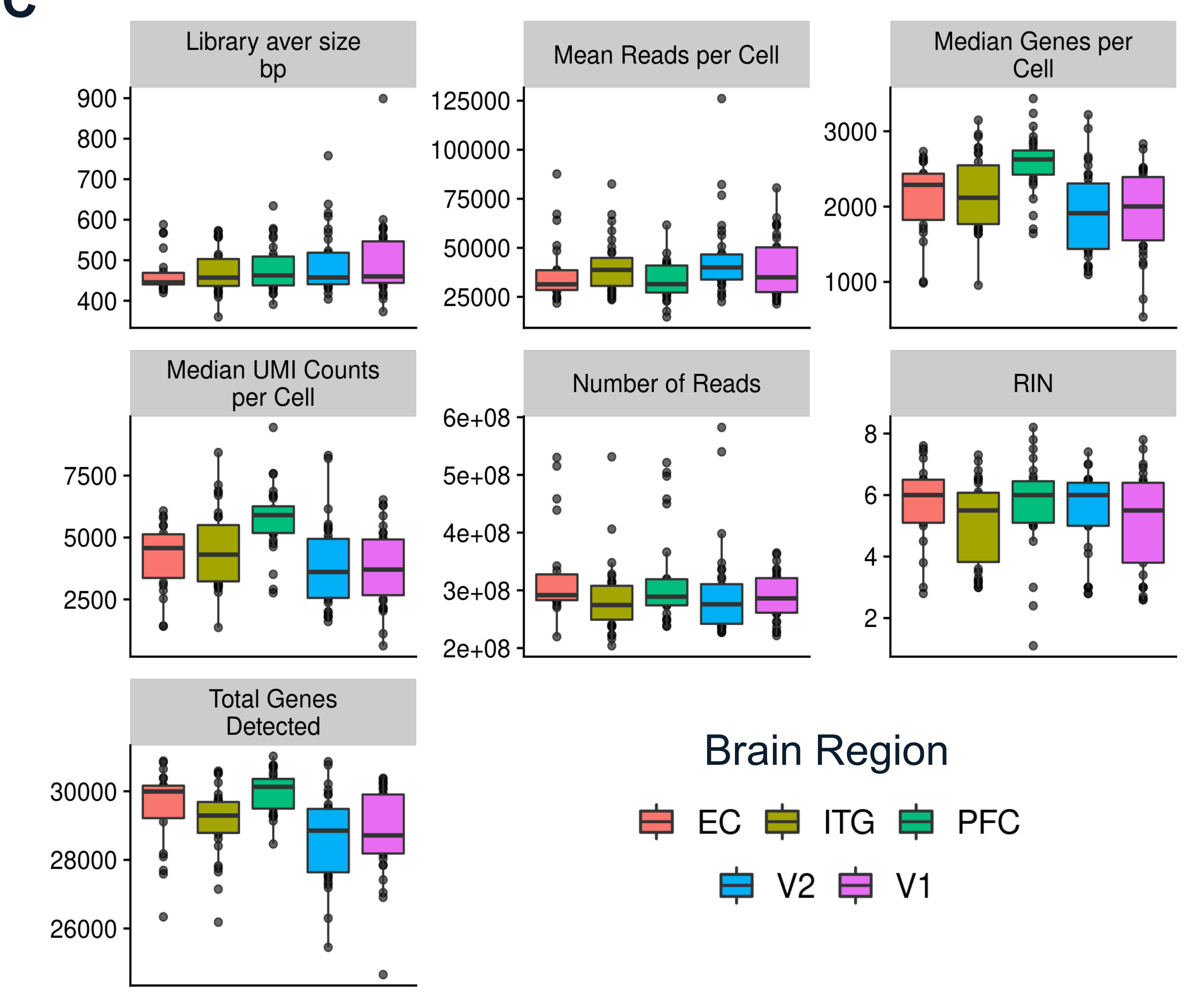


**D**

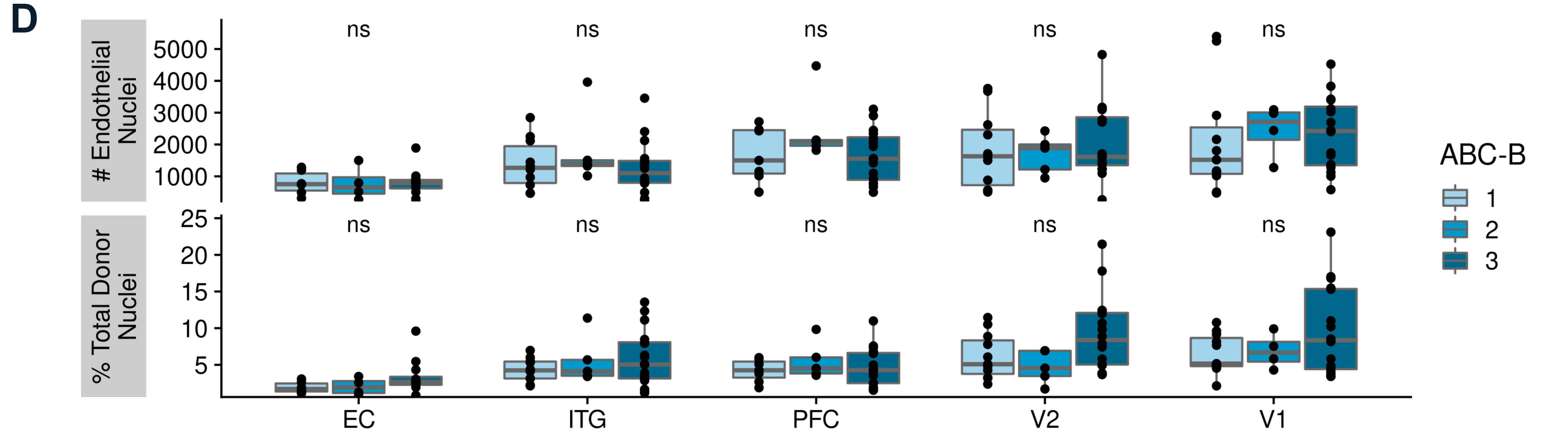

Figure S3

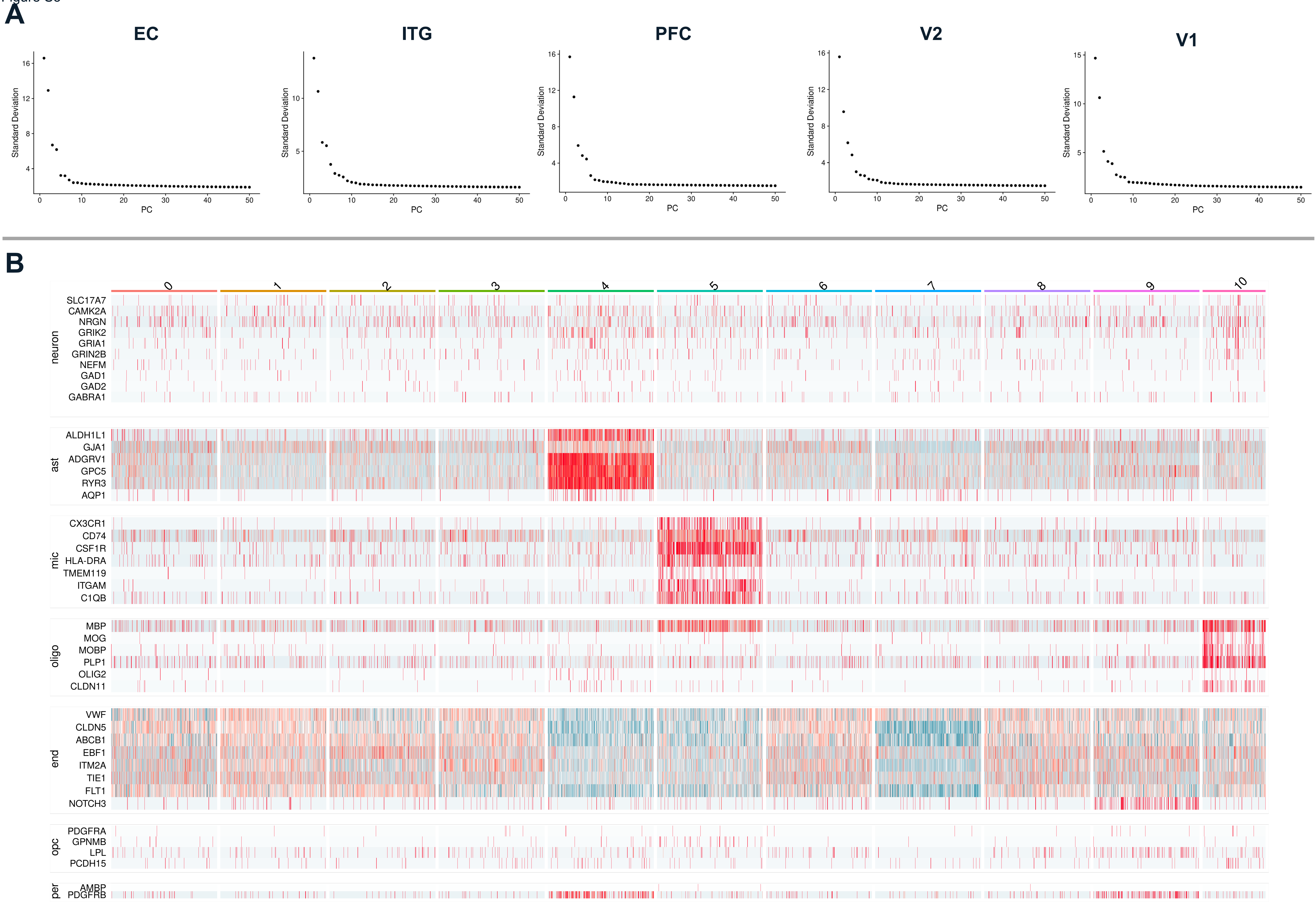
A
EC
ITG
PFC
V2
V1
Standard Deviation
PC
B
0
1
2
3
4
5
6
7
8
9
10
neuron
SLC17A7
CAMK2A
NRGN
GRIK2
GRIA1
GRIN2B
NEFM
GAD1
GAD2
GABRA1
ast
ALDH1L1
GJA1
ADGRV1
GPC5
RYR3
AQP1
mic
CX3CR1
CD74
CSF1R
HLA-DRA
TMEM119
ITGAM
C1QB
oligo
MBP
MOG
MOBP
PLP1
OLIG2
CLDN11
end
VWF
CLDN5
ABCB1
EBF1
ITM2A
TIE1
FLT1
NOTCH3
opc
PDGFRA
GPNMB
LPL
PCDH15
per
AMBP
PDGFRB

Figure S4

**A**

| Segment | # Nuclei | Median # Genes |
|---|---|---|
| Arterial | 5,046 | 2,737.5 |
| Capillary | 41,517 | 2,893 |
| Venous | 5,023 | 3,647 |

**B**

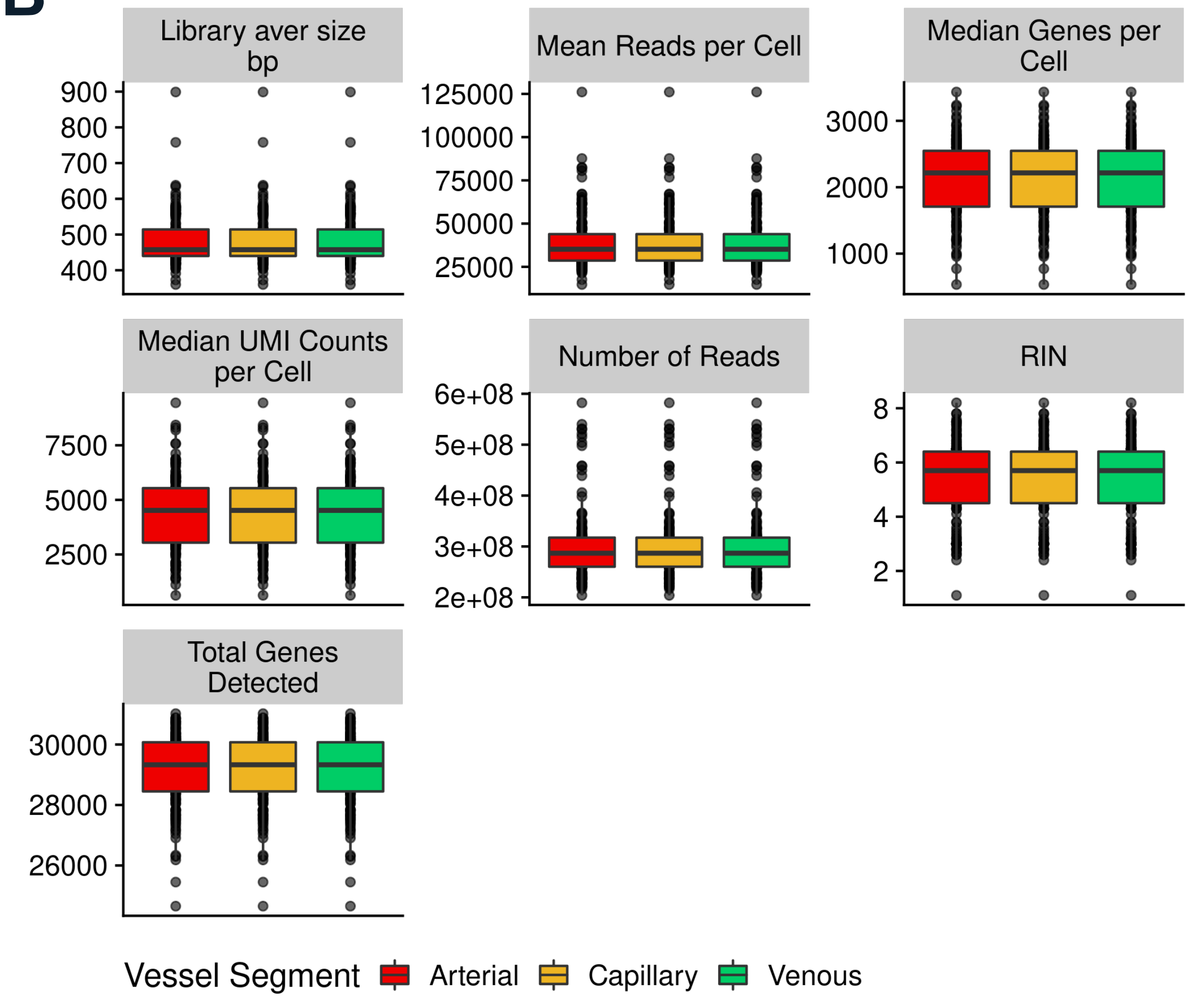


**C**

Vessel-Specific AD DEG counts

| | Arterial | Capillary | Venous |
|---|---|---|---|
| Up | 5 | 474 | 17 |
| Down | 4 | 505 | 12 |

Figure S5

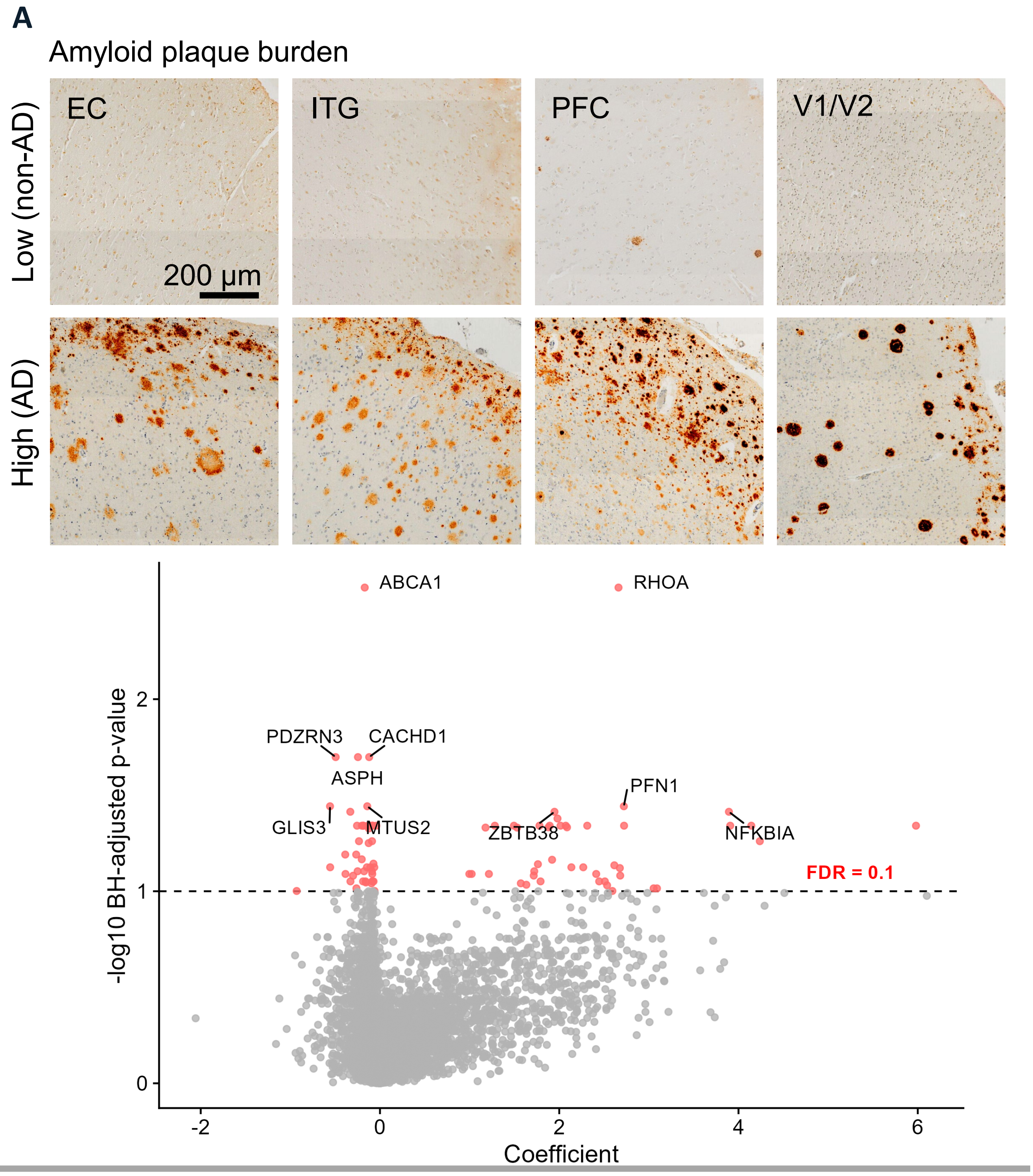
A
Amyloid plaque burden
Low (non-AD)
High (AD)
EC
ITG
PFC
V1/V2
200 µm
-log10 BH-adjusted p-value
Coefficient
ABCA1
RHOA
PDZRN3
CACHD1
ASPH
PFN1
GLIS3
MTUS2
ZBTB38
NFKBIA
FDR = 0.1
-2
0
2
4
6
1
2

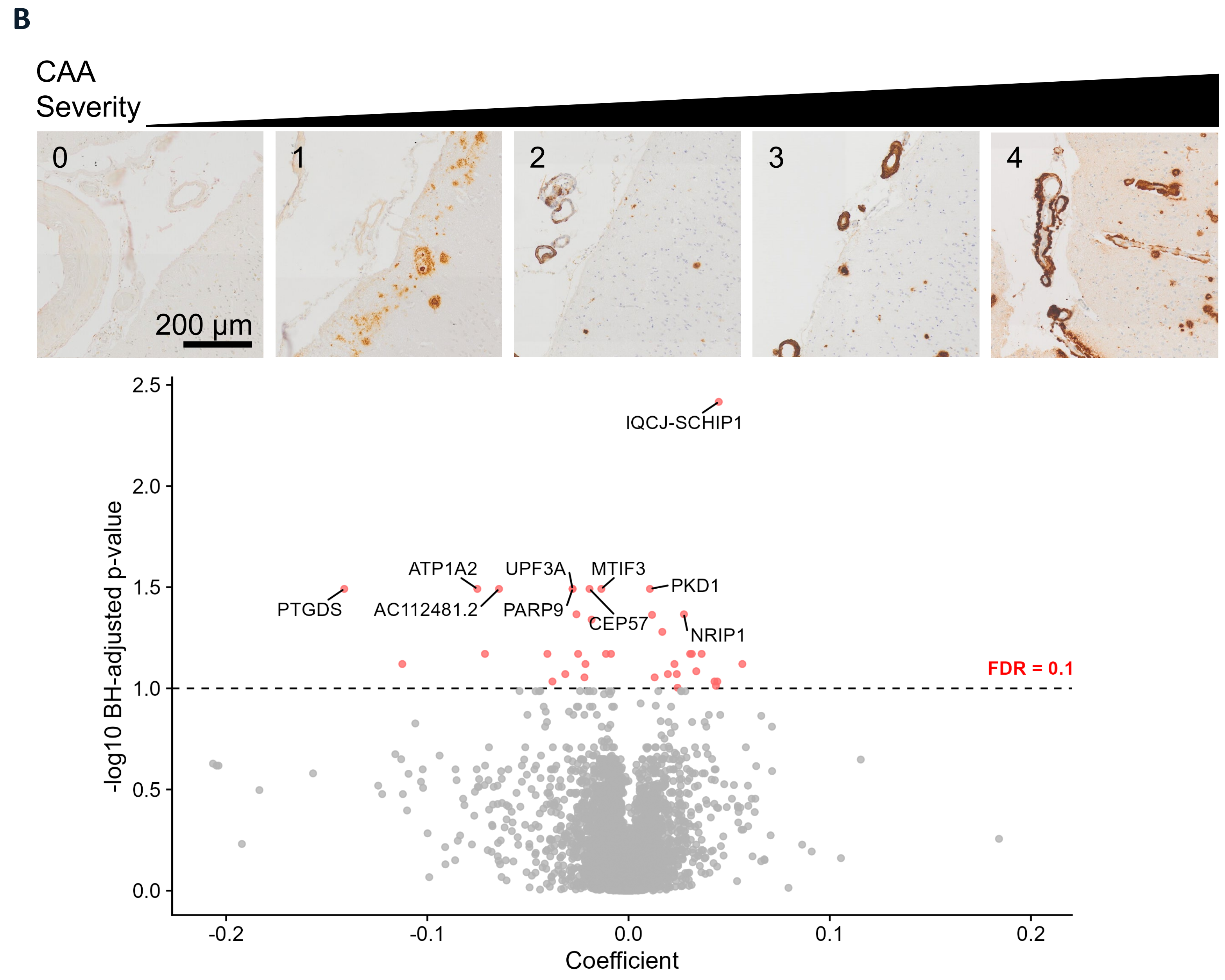
B
CAA
Severity
0
1
2
3
4
200 µm
-log10 BH-adjusted p-value
Coefficient
IQCJ-SCHIP1
ATP1A2
UPF3A
MTIF3
PKD1
PTGDS
AC112481.2
PARP9
CEP57
NRIP1
FDR = 0.1
2.5
2.0
1.5
1.0
0.5
0.0
-0.2
-0.1
0.1
0.2
